# Zinc Tolerance Through Glutathione Import Strikes a Fine Balance Between Protection and Damage in *Streptococcus mutans*

**DOI:** 10.64898/2026.06.01.728743

**Authors:** Miranda C. Carter, Edroyal Womack, Malak Khatib, Alexandra M. Peterson, Irene S. Saengpet, José A. Lemos

## Abstract

Recently, our group showed that the dental pathogen *Streptococcus mutans* is inherently more tolerant to high zinc stress than other streptococci, a phenotype associated with the presence of a P-type ATPase exporter named ZccE, virtually unique to *S. mutans*. In addition to *zccE*, a previous transcriptome analysis revealed that *S. mutans* upregulates genes involved in glutathione uptake during initial exposure to zinc stress. Glutathione, a major supplier of organic sulfur that also plays key roles in antioxidant defense and xenobiotic detoxification, forms coordination complexes with a variety of metals, including zinc, thereby functioning as a buffer that protects cells from metal intoxication. To investigate the contribution of glutathione zinc tolerance in *S. mutans*, the *gshT* gene, which encodes the substrate-binding subunit of a glutathione transporter, was deleted in both the parent and Δ*zccE* strains and the ability of these mutants to overcome zinc stress through intracellular glutathione accumulation determined. Targeted metabolomics revealed that *S. mutans* accumulates glutathione in a GshT-dependent manner following zinc stress, a response that was strikingly amplified in the Δ*zccE* strain. Although glutathione supplementation had a minimal and non-significant impact on growth of either parent or mutant strains in sub-inhibitory zinc concentrations, the Δ*gshT* strain exhibited increased zinc sensitivity in a plate-based assay. However, the Δ*zccE*Δ*gshT* mutant displayed enhanced zinc tolerance compared to the Δ*zccE* single mutant. While glutathione alone did not alter zinc levels in the UA159 or Δ*zccE* strains, the combination of zinc and glutathione nearly doubled intracellular zinc levels in Δ*zccE* compared to cells grown in zinc only. We conclude that while glutathione may play a minor role in *S. mutans* zinc tolerance, uncontrolled glutathione uptake observed in Δ*zccE* facilitates zinc entry, as glutathione:Zn^2+^ complexes inadvertently promote zinc intoxication via a Trojan horse mechanism.

## INTRODUCTION

The World Health Organization (WHO) describes dental caries as the most common human condition globally, affecting over 2.5 billion people annually and imposing an economic burden of nearly 250 billion USD (1). Although multifactorial, dental caries onset is strongly linked to increased *Streptococcus mutans* presence in the supragingival biofilm (2,3). While typically present at low levels or absent in healthy dental biofilms, *S. mutans* thrives in environments with high sucrose availability, which is utilized to produce large amounts of extracellular glucans that facilitate the formation of robust biofilms and concurrently lower the environmental pH (2,4). These abilities enable *S. mutans* to create an acidic biofilm environment that is unhospitable to oral commensals, decreasing microbial diversity while selecting for highly aciduric, caries-associated microorganisms (4,5).

As the second most abundant trace metal in the human body and both an essential and toxic mineral, zinc is deeply involved in host-pathogen interactions (6–8). Specifically, during infections, as part of the host innate immunity response, free zinc is actively sequestered by calprotectin, a heterodimer of the S100A8 and S100A9 proteins that is profusely secreted by neutrophils and other immune cells starving the invading pathogen of this essential trace metal (6). Simultaneously, zinc is mobilized within macrophages and other phagocytic cells, facilitating the killing of pathogens entrapped in phagosomes through disruption of critical metabolic processes (9,10). As a toxic agent, zinc’s high binding affinity for protein ligands, indicated by its placement just below copper in the Irving-Williams series of metal stability, enables it to inadvertently bind proteins that normally require other metal cofactors, forming stable interactions that disrupt their function, a process known as protein mismetallation (11,12). In an effort to maintain intracellular zinc homeostasis in the face of zinc starvation or excess, bacteria have evolved high-affinity import and export systems, as well as auxiliary mechanisms that help them adapt to fluctuating environmental zinc conditions (9).

For centuries, the antimicrobial and anti-inflammatory properties of zinc combined with its relatively low toxicity to mammalian cells has led to the utilization of zinc as a therapeutic agent. Human conditions including the common cold and other respiratory infections, wound healing, and the reduction of duration and severity of pediatric diarrhea have all shown improvement with zinc supplementation (13–15). Moreover, in oral health, zinc salts have frequently been incorporated into oral care products to treat gingivitis and halitosis and, arguably, aid in the control of dental plaque build-up (16–18).

Given the strong presence of zinc in oral care formulations, we have recently begun to investigate the effects of supraphysiological zinc on *S. mutans* biology. We discovered that, despite lacking the canonical zinc exporter CzcD typically found in members of the *Streptococcus* genus, *S. mutans* is far more tolerant to zinc stress than all other oral streptococci tested (19,20). This unexpected observation was accompanied by the discovery of ZccE, a multi-metal exporter from the P_1B_ -type ATPase family, unique to *S. mutans*, that was directly responsible for *S. mutans’* high zinc tolerance (19). In addition to *zccE*, analysis of the zinc stress transcriptome of *S. mutans* revealed additional pathways that may contribute to zinc tolerance (19). Among the genes of interest upregulated during an early stage of high zinc stress were genes associated with glutathione transport, utilization, and regeneration (19). A multifunctional tripeptide of cysteine, glutamate, and glycine, glutathione is an abundant low molecular weight thiol, existing in the reduced (GSH) and oxidized (GSSG) forms. Recognized for its antioxidant properties and major role in redox homeostasis, glutathione fulfills multiple critical functions in oxidative stress defense, including direct scavenging of reactive oxygen species (ROS), serving as a cofactor for glutathione peroxidases, and preserving protein thiol groups to maintain proper structure and protein stability (21–26). Glutathione also mediates the detoxification of xenobiotics, including toxic heavy metals such as cadmium, lead and mercury, by forming GSH complexes that are more efficiently eliminated from the cell. Finally, GSH interacts with trace biometals such as copper and zinc serving as an intracellular buffer that protects cells from metal-induced oxidative stress and direct metal intoxication. In *E. coli*, glutathione was shown contribute to tolerance against the toxic metals cadmium and chromate, whereas cells that cannot synthesize or import glutathione display heightened sensitivity to copper and zinc, especially in the absence of the CopA and ZntA exporters, respectively (27). In streptococci, glutathione has been shown to mediates copper tolerance in *S. pyogenes,* and cadmium, copper, and zinc tolerance in *S. pneumoniae* (21,28).

Considering the established intracellular metal buffering capacity of glutathione and our recent finding that transcription of the *S. mutans* glutathione import genes (*tcyABC* and *gshT*) (29) is induced shortly after high zinc stress (19), here we investigated the role of glutathione in *S. mutans* zinc tolerance. We report that while increases in intracellular glutathione pools during stress play only a minor role in *S. mutans* zinc tolerance, dysregulated uptake causes glutathione to function as a ’Trojan horse’ facilitating zinc entry and exacerbating zinc intoxication. Specifically, uncontrolled glutathione import, observed in the Δ*zccE* zinc export mutant as a potential compensatory mechanism, enhanced zinc intoxication by inadvertently promoting the uptake of GSH:Zn²⁺ complexes.

## METHODS

### Strains and growth-based experiments

The strains used in this study are listed in Table 1. All strains were routinely grown in brain heart infusion (BHI) broth or BHI agar plates at 37°C in 5% CO_2_. When appropriate, media was supplemented with 1 mg ml^-1^ spectinomycin, 1 mg ml^-1^ kanamycin, or 10 µg ml^-1^ erythromycin. To generate growth curves with *S. mutans*, overnight cultures of serotype *c* strain UA159 and derivatives were sub-cultured in fresh media, grown to mid-log phase (OD_600_ ∼ 0.4), diluted 1:50 into the desired condition, and their growth monitored using an automated growth plate reader (Bioscreen C; Oy Growth Curves Ab) set to 37°C with each well overlayed with sterile mineral oil to minimize generation of reactive oxygen species.

**Table 1.**
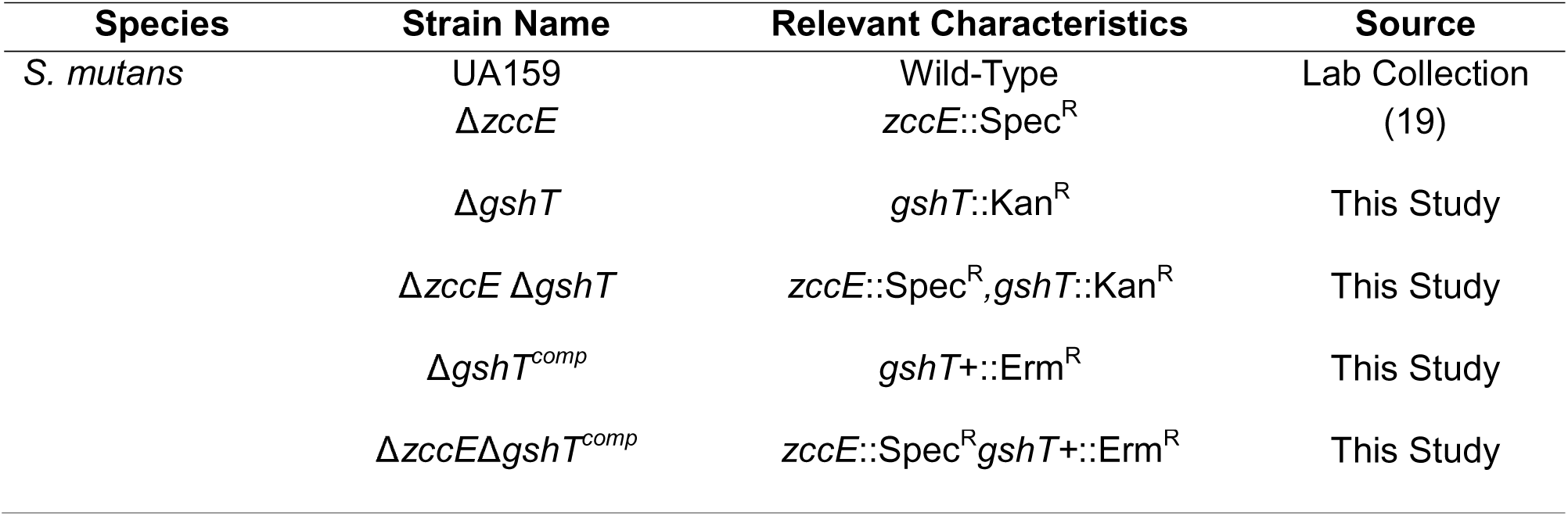
Strains used in the study.

For plate titration assays, mid-log phase (OD_600_ ∼0.4) cultures were serially diluted in phosphate-buffered saline (PBS) and 5 µL of 5- or 10-fold dilutions spotted on BHI agar with or without metal supplementation. Plates were incubated for 48 hours at 37°C in 5% CO_2_ before images were captured. To assess strain viability in the presence of zinc and/or glutathione, mid-log grown cultures were exposed to 1 mM ZnSO_4_, 100 µM glutathione, or both for 90 minutes, the cell pellets obtained by centrifugation, washed in sterile PBS, serially diluted and 10-fold dilutions spotted on BHI agar. Plates were incubated for 24 hours at 37°C in 5% CO_2_ before images were captured.

### Construction of mutant and complemented strains in *S. mutans*

Strains of *S. mutans* lacking *gshT* were constructed using a PCR ligation mutagenesis approach (30). Briefly, DNA fragments flanking the target gene were amplified by PCR using the primers listed in Table S1 and ligated to a non-polar kanamycin resistance cassette. The ligated fragment was used to transform the UA159 and Δ*zccE* strains using an established protocol (31). Mutant strains were isolated on BHI plates supplemented with kanamycin, and *gshT* deletion on each strain confirmed by sequencing amplicons containing the antibiotic cassette insertion site and flanking region. To generate complemented strains, the *gshT* coding sequence and flanking region was amplified from UA159 genomic DNA, and an erythromycin resistance cassette inserted upstream of the native *gshT* promoter using Gibson Assembly (32). This fragment was used to transform the Δ*gshT* and Δ*zccE*Δ*gshT* strains in the presence of 1 mM competence stimulating peptide (CSP) (31). Strains were selected on BHI agar supplemented with erythromycin and *gshT* re-integration verified via Sanger sequencing.

### Quantitative real time PCR

*S. mutans* UA159 and Δ*zccE* were grown in BHI to mid-log phase and treated with 1 mM ZnSO_4_ for 15 min. The cell pellets were collected by centrifugation, resuspended in RNA Protect reagent (Qiagen) and stored at -80°C prior to RNA extraction. Total RNA was isolated from homogenized *S. mutans* lysate using the acid-phenol-chloroform extraction methods as previously described (33). Total RNA was treated with Ambion DNase I (ThermoFisher) for 30 minutes at 37°C, followed by another purification step using the RNeasy Mini Kit (Qiagen) and treatment with the Turbo DNA-free kit (Invitrogen). Reverse transcription and qRT-PCR reactions were carried out with the High-Capacity cDNA Reverse transcription Kit (Applied Biosystems) and the iTaq Universal SYBR Green Supermix (Bio Rad), using the primers listed in Table S1. Expression of *gshT, tcyA,* and *gshAB* was quantified via ΔΔC_T_, where C_T_ represents the threshold cycle using *gyrA* as the housekeeping gene (34).

### Inductively coupled mass spectrometry

Intracellular zinc and manganese concentrations of *S. mutans* strains before and after exposure to ZnSO_4_, glutathione, or both were determined via inductively coupled plasma mass spectrometry (ICP-MS) performed at the University of Florida Center of Environmental and Human Toxicology Analytical Toxicology core laboratory. Briefly, cells were harvested by centrifugation at 4°C for 15 min at 4,000 rpm, the cell pellets resuspended in 1 ml of 50% (v/v) HNO_3,_ and 0.2 ml 30% (v/v) H_2_O_2_ and heated at 100°C for 1 hour. Once cool, 0.2 ml of 30% (v/v) H_2_O_2_ were added to each sample, and samples were brought to a total volume of 16 ml with ultrapure water. Intracellular metal content determined using a 7900 ICP Mass Spectrometer (Agilent) in helium gas mode to minimize polyatomic interferences. Metal concentrations were normalized to total protein content determined by the bicinchoninic acid (BCA) assay (Pierce).

### Intracellular glutathione quantification

Glutathione levels were determined using a high-performance mass spectrometer approach developed in conjunction with the University of Florida Mass Spectrometry Research and Education Center (MSREC). Briefly, samples were grown in BHI to mid-log phase (OD_600_ ∼0.4, control conditions) before being exposed to zinc (2 mM ZnSO_4_ for UA159 and Δ*gshT* strains and 1 mM ZnSO_4_ for Δ*zccE*), 100 μM glutathione, or both for 45 or 90 minutes. Cells were collected by centrifugation (7,000 rpm for 5 minutes at 4°C), washed twice in sterile PBS, dried in a SpeedVac, and stored at -20°C. Metabolite extraction was performed by resuspending cell pellets in 300 μl of cold 90% methanol containing 5 mM dithiothreitol (DTT), sonicated in ice water for 10 minutes and frozen at -20°C for 1 hr. Then, samples were centrifuged at 14,000 rpm for 5 min, and the supernatant stored at -20°C. This process was repeated with cold 100% methanol containing 5 mM DTT, and cold 90% acetonitrile containing 5 mM DTT. Supernatant from each extraction was pooled and dried using a Speed Vac before storage at -20°C until reconstitution in 100 µl H_2_O with 5 mM DTT. Mass spectrometry (MS) analysis was performed on a TIMS-TOF Pro II TIMS-qTOP MS with a ThermoScientific Hypersil GOLD PFP column under a 20 min gradient using water with ammonium formate and methanol as mobile phases, both containing formic acid. For data normalization, 1 ml culture aliquots were washed, serially diluted in PBS, and plated on BHI agar to determine colony forming units per ml.

### Statistical analysis

Data obtained from this study were analyzed using R (*version 4.5.0,* http://www.r-project.org) in RStudio (*version 2026.1.0.392; Posit Software, PBC, Boston, MA.* http://www.posit.co/) as described in figure legends. R packages used for visualization and data analysis are listed in Supplemental Table S2. Unless stated otherwise, only comparisons with significant differences (p-value < 0.05) indicated with its corresponding asterisks were noted in the figure legends.

## RESULTS

### Inactivation of *gshT* impairs *S. mutans* growth kinetics and metal tolerance

In the first set of experiments, we revisited the transcriptome results using quantitative real-time PCR to obtain the transcriptional profile of *gshT* and *tcyA* in response to zinc stress in both the UA159 (parent) and Δ*zccE* (zinc export mutant) strains. To adjust for the high zinc sensitivity and loss of viability of the Δ*zccE* strain (19), we exposed cultures to 1 mM ZnSO_4_ for 15 minutes instead of 4 mM ZnSO_4_ that was used in the original study. Zinc exposure in UA159 led to a moderate upregulation of *gshT* (∼2-fold) but no changes in *tcyA* expression. On the other hand, the upregulation of *gshT* (4-fold) was substantially greater in Δ*zccE* with *tcyA* (3-fold) also showing a significant increase. These results support that *S. mutans* upregulates glutathione import in response to high zinc stress while also indicating that increased glutathione uptake may act as a compensatory response to the loss of zinc export activity (Fig. 1). Upon confirming a potential association between glutathione import with zinc stress survival, we next asked whether supplementation of the growth medium with glutathione could improve growth of UA159 under a sub-inhibitory concentration of ZnSO_4_. While glutathione concentrations at or above 200 µM reduced final growth yields, 100 µM glutathione supplementation had no noticeable effect on growth in the absence of zinc stress (Fig. S1A). We then tested whether the addition of 100 µM glutathione to BHI improved growth in the presence of a sub-inhibitory concentration of zinc, observing a very modest increase in growth yield after media supplementation (Fig. S1B).

**Fig. 1.**
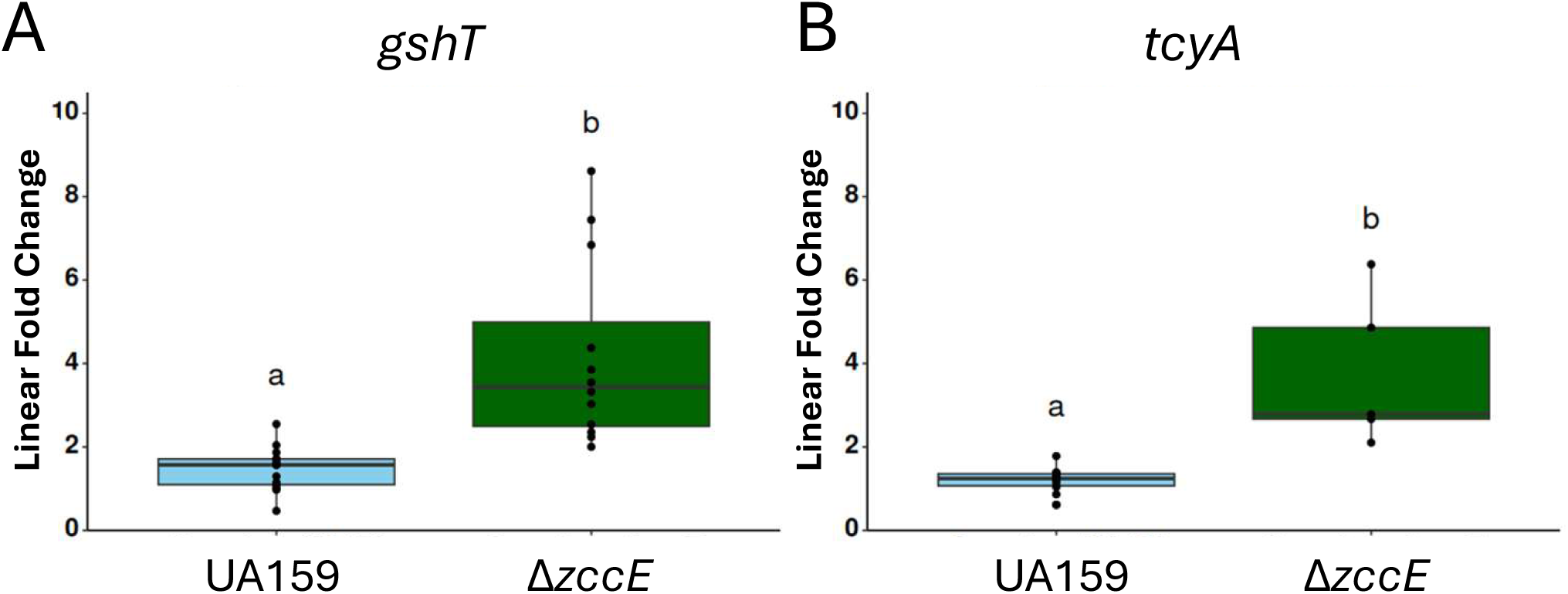
Glutathione import genes are induced by zinc stress. qRT PCR was performed using ΔΔCt normalization to *gyrA.* Strains (UA159 and Δ*zccE*) were grown to mid-log phase in BHI and exposed to 1 mM ZnSO_4_ for 15 minutes. Error bars represent standard deviation. Differing letters indicate significant difference (p<0.05) between groups according to Wilcox Rank Sum (p< 0.05) test that was used due to non-normality.

To further investigate the role of glutathione in zinc tolerance, we generated a *gshT* deletion mutant (Δ*gshT*) to assess its growth characteristics under zinc stress. Given that the ZccE exporter is primarily responsible for zinc tolerance (19), we also generated a Δ*zccE*Δ*gshT* double mutant to probe for a potential epistatic effect. In plain BHI, that contains low (∼10 µM) zinc concentrations(35), inactivation of *gshT* in the UA159 parent or Δ*zccE* strain backgrounds greatly reduced final growth yields, a phenotype that was rescued by genetic complementation of the *gshT* gene (Fig. S2A-B). For unclear reasons, this growth defect became less pronounced in the presence of sub-inhibitory zinc concentration (Fig. S2C-D). Because the pronounced growth defect of Δ*gshT* strains in broth creates a confounding factor for interpretation of growth kinetic assays, we instead investigated the linkage between glutathione and zinc tolerance using a plate titration assay that assesses endpoint growth, with all strains showing comparable growth on BHI plates after 48 hours of incubation (Fig. 2A-B). On plates containing zinc, growth of the Δ*gshT* strain was slightly impaired when compared to the parent strain UA159, providing direct evidence of glutathione’s role in zinc stress mitigation (Fig. 2A). Unexpectedly, this phenotype was reversed in the Δ*zccE* background with the Δ*zccE*Δ*gshT* double mutant growing better on zinc-containing plates than the single mutant (Fig. 2B). Because ZccE mediates tolerance to other trace metals in addition to zinc (19), we also evaluated the ability of the *gshT* mutants to grow in the presence of cadmium and cobalt. This time, inactivation of *gshT* increased cadmium sensitivity in UA159 and even more so in Δ*zccE* while increased cobalt sensitivity was observed in the Δ*zccE* background (Fig. 2A-B).

**Fig. 2.**
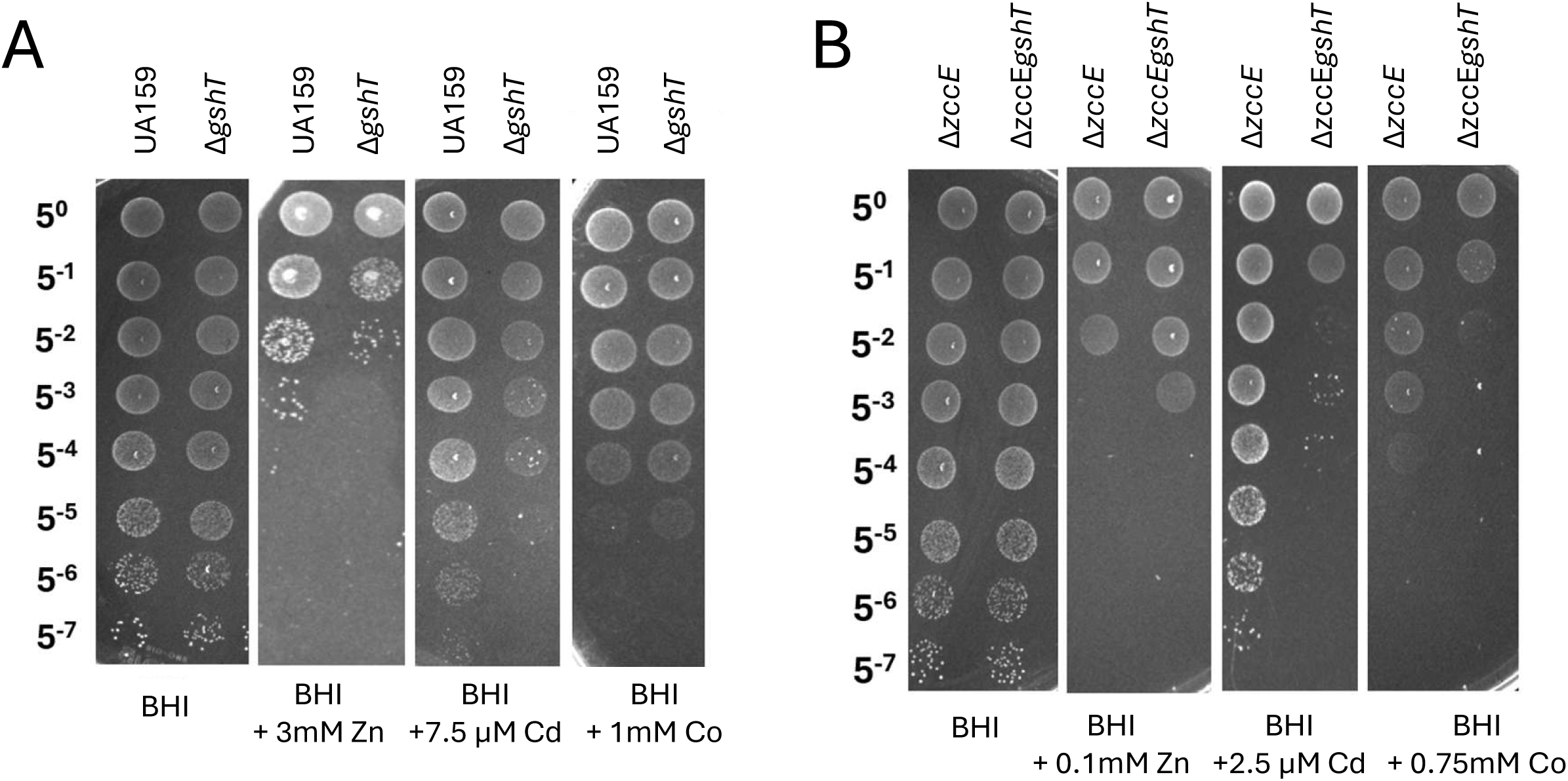
Inactivation of *gshT* impacts metal tolerance. Spot test of *S. mutans* UA159 and Δ*gshT* (A) or Δ*zccE* and Δ*zccE*Δ*gshT* (B) on BHI plates with or without sub-inhibitory concentrations of zinc (Zn), cadmium (Cd) or cobalt (Co). The different concentrations of each metal used for UA159 and Δ*zccE* background strains are indicated in the figure. Images were taken after 48 hours incubation at 37°C in a 5% CO_2_ and are representative of at least 3 independent experiments.

### Glutathione uptake greatly increases under zinc stress

Next, we sought to determine intracellular glutathione levels in the UA159, Δ*zccE*, and Δ*gshT* strains before and after zinc stress. For this, strains were grown with or without supplementation of 100 µM glutathione and exposed to zinc stress (2 mM ZnSO_4_ for both UA159 and Δ*gshT* and 1 mM ZnSO_4_ for Δ*zccE*) for up to 90 minutes. Regardless of the growth condition, glutathione was not detected in the Δ*gshT* strain (Fig. 3A), indicating that *S. mutans* UA159 is completely dependent on GshT-mediated import to maintain detectable levels of glutathione and, most likely, cannot synthesize glutathione. However, intracellular glutathione levels nearly tripled in the parent strain after zinc stress, increasing even further (∼5-fold) when the medium was supplemented with glutathione and then subjected to zinc stress (Fig. 3B). Furthermore, zinc-induced import/accumulation of glutathione was much greater in the Δ*zccE* strain that displayed intrinsic higher levels of glutathione in the absence of zinc stress that tripled after zinc stress and increased by nearly 12-fold after glutathione supplementation (Fig. 3C). These findings validate the transcriptional data indicating that *S. mutans* imports glutathione to mitigate zinc stress while also supporting our initial hypothesis that the Δ*zccE* strain increases glutathione import as a compensatory mechanism to overcome zinc stress.

**Fig. 3.**
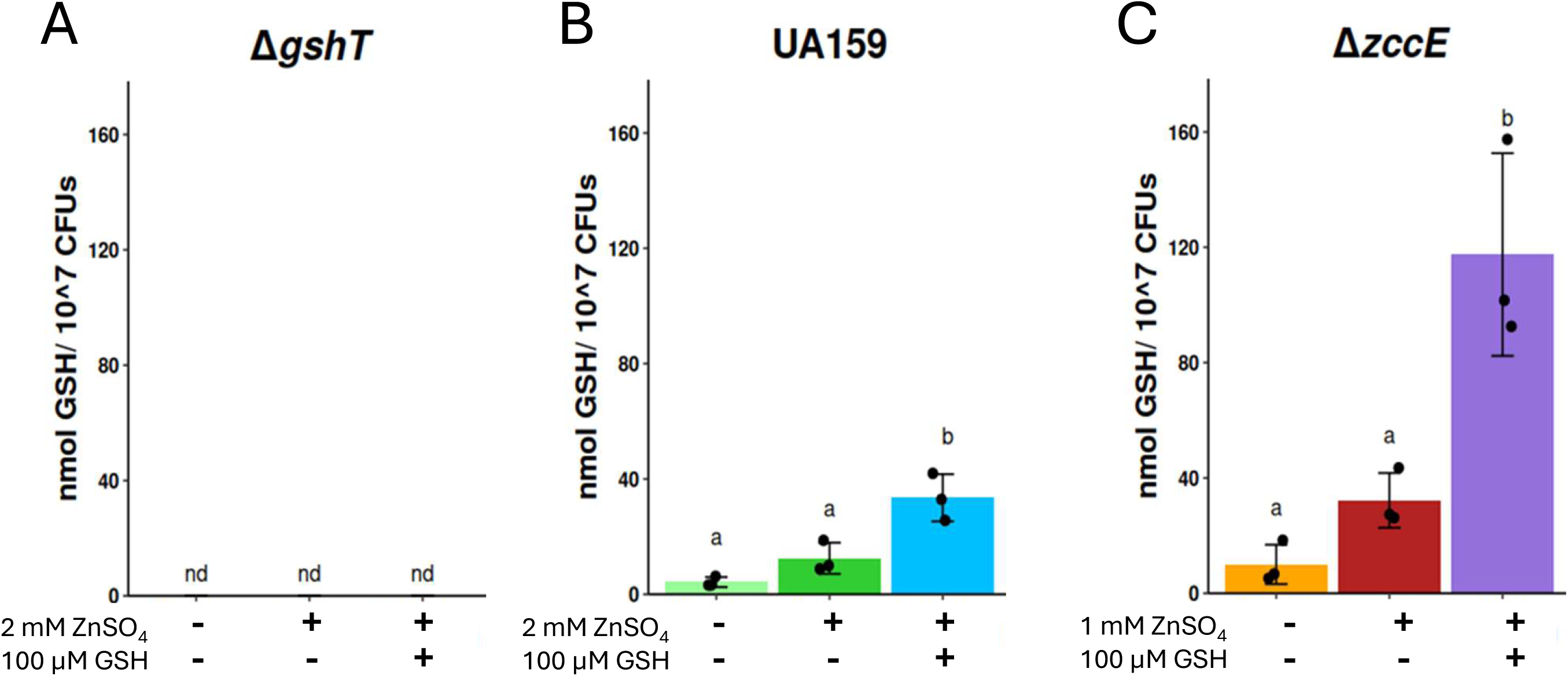
Zinc stress triggers glutathione uptake. Cultures of UA159, Δ*gshT* and Δ*zccE* were grown to mid-log phase and exposed to zinc stress (2 mM ZnSO_4_ for UA159 and Δ*gshT* strains and 1 mM ZnSO_4_ for Δ*zccE*), 100 µM glutathione (GSH), or both for 90 minutes. Intracellular glutathione was determined by mass spectrometry and the data normalized by CFUs. Error bars represent standard deviation. Differing letters indicate significant difference (p<0.05), while groups denoted with the same letter are not significantly different (p> 0.05) between groups according to two-way ANOVA with Bonferroni’s correction for multiple comparisons. (nd) Not detected.

### Intracellular zinc quantification suggests a Trojan horse mechanism

Beyond the evidence that increased glutathione uptake helps *S. mutans* cope with metal stress, albeit playing a small role, our results also suggest that uncontrolled glutathione import is harmful to the bacterium under high zinc conditions. Because inactivation of *gshT* improved zinc tolerance of the Δ*zccE* strain (Fig. 2B), we suspected that glutathione may inadvertently function as a ’Trojan horse’ facilitating zinc entry through GshT-mediated import of GSH:Zn complexes. While less likely, another possibility is that the uncontrolled uptake and accumulation of glutathione alone is toxic given that supplementation of the growth medium with high concentrations of glutathione inhibited bacterial growth (Fig. S1A). To investigate the first possibility, we determined intracellular zinc concentrations in UA159, Δ*gshT*, Δ*zccE,* and Δ*zccE*Δ*gshT* strains grown to mid-log phase in BHI (control) or exposed for 90 minutes to 1 mM ZnSO_4_, 100 µM glutathione, or both. Though glutathione alone did not affect intracellular zinc levels in either UA159 or Δ*gshT*, zinc treatment, with or without glutathione supplementation, resulted in a slight, non-significant increase in intracellular zinc (Fig. 4A). Glutathione supplementation alone also did not alter zinc levels in the Δ*zccE* or Δ*zccE*Δ*gshT* strains but, as shown previously (19), zinc treatment led to a very substantial elevation (∼12-fold) in Δ*zccE* that, remarkably, nearly doubled after glutathione supplementation (Fig. 4A). Finally, zinc accumulation was significantly less pronounced in the Δ*zccE*Δ*gshT* strain when compared to the Δ*zccE* single mutant, with the addition of glutathione having no further impact. As expected based on the well-established inverse relationship between manganese and zinc (36–38), intracellular manganese levels were significantly reduced in the Δ*zccE* and Δ*zccE*Δ*gshT* strains (Fig. 4B). While the reasons are unclear, glutathione supplementation, in the absence of zinc stress, significantly reduced intracellular manganese in Δ*zccE* and even more so in Δ*zccE*Δ*gshT* (Fig. 4B).

**Fig. 4.**
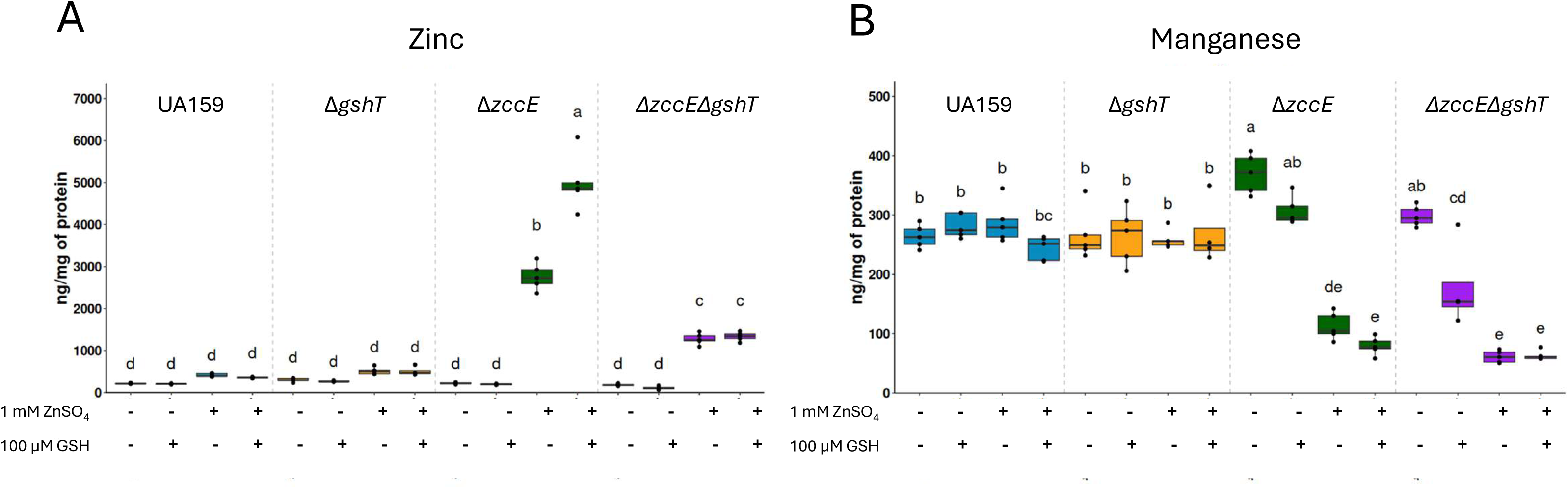
Uncontrolled glutathione import via GshT facilitates zinc entry. Mid-log grown BHI cultures of UA159, Δ*zccE,* Δ*gshT,* and Δ*zccE* Δ*gshT* were treated with 1 mM ZnSO_4_, 100 µM glutathione (GSH), or both, for 90 minutes and intracellular zinc (A) and manganese (B) quantified via ICP-MS. Error bars represent standard deviation. Differing letters indicate significant difference (p<0.05), while groups denoted with the same letter are not significantly different (p> 0.05) between groups according to two-way ANOVA with Tukey’s multiple comparison test.

To support the proposed Trojan Horse mechanism, we sought additional evidence that the increased glutathione uptake observed in the Δ*zccE* strain triggers inadvertent cell death due to zinc intoxication by incorporation of GSH:Zn complexes. Here, we used the same experimental setup as with ICP-MS analysis, in which mid-log cultures were treated for 90 minutes with 1 mM ZnSO_4_, 100 µM glutathione, or both, followed by plating to determine cell viability. We found that while zinc exposure alone had no impact on the viability of any of the strains, co-treatment with glutathione and zinc diminished viability of Δ*zccE,* a phenotype that was rescued with inactivation of *gshT* (Fig. 5).

**Fig. 5.**
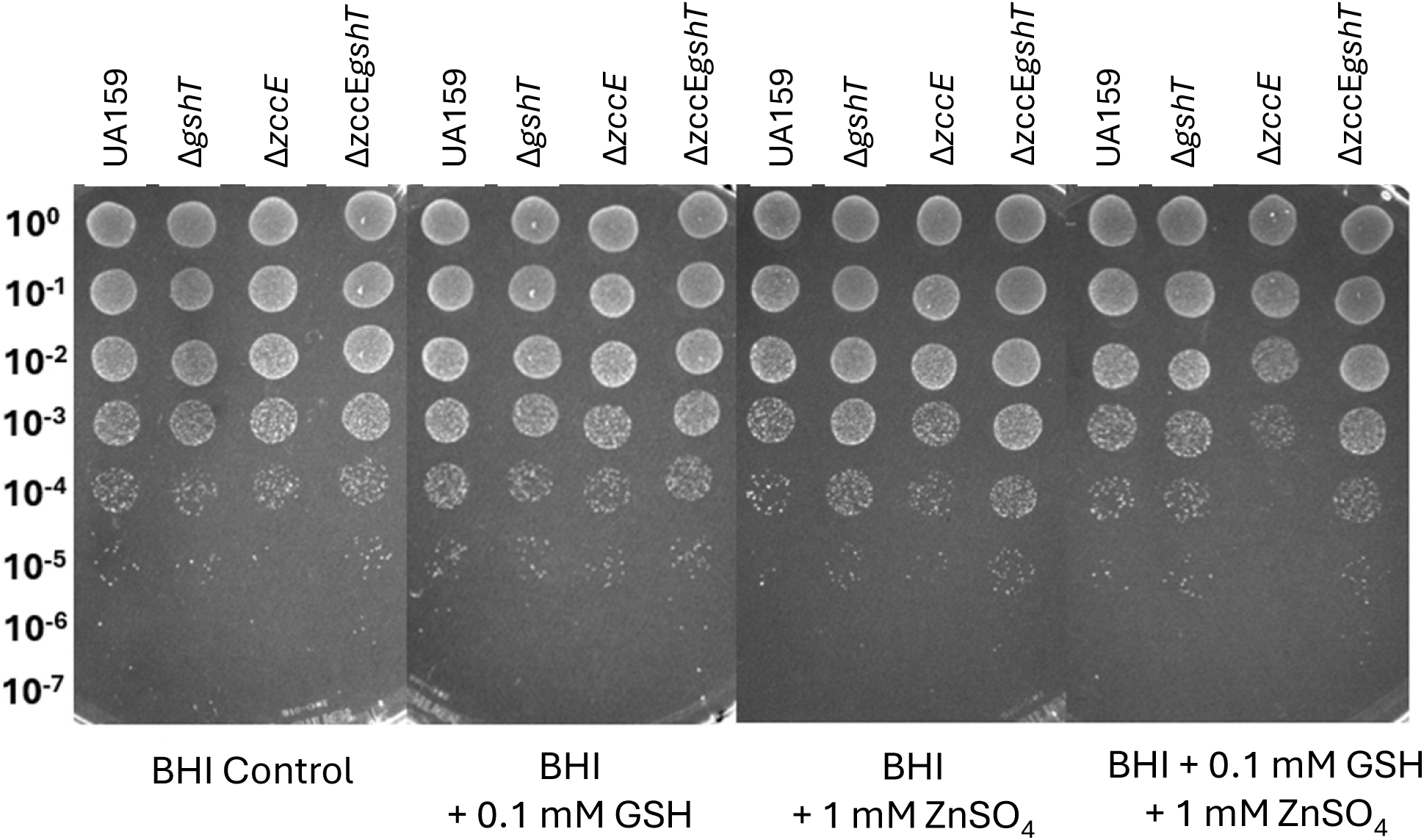
Uncontrolled uptake of GSH-Zn^2+^ complex is toxic to the Δ*zccE* strain. Mid-log grown BHI cultures of UA159, Δ*zccE,* Δ*gshT,* and Δ*zccE* Δ*gshT* were treated with 1 mM ZnSO_4_, 0.1 mM GSH, or both, for 90 minutes and cell viability determined by plating serial dilutions on BHI agar.

## DISCUSSION

In *S. mutans*, zinc export is mediated by ZccE, a P-type ATPase virtually unique to the species, which grants a substantially greater zinc tolerance in comparison to other streptococci that utilize the widely distributed CDF-type exporter CzcD to cope with high zinc stress (19,39,40). In our previous work, we identified several genes, apart from *zccE*, that were upregulated during zinc stress, which included genes associated with glutathione uptake, utilization, and regeneration (19). As the major thiol species in prokaryotes and eukaryotes, glutathione serves a variety of roles including detoxification of metals by sequestration/buffering and efflux through export of metal:GSH complexes (41). In this report, we explored the potential of glutathione to function as an intracellular buffer potentially serving as an auxiliary mechanism to ZccE-mediated zinc export.

Prior studies have characterized the mechanisms of glutathione uptake in *S. mutans* revealing that GshT, a type 3 solute binding protein, partners with the TcyB permease and TcyC ATPase from the TcyABC L-cystine transporter to import exogenous glutathione (29,42). Here, inactivation of *gshT* resulted in no detectable intracellular glutathione indicating that *S. mutans* relies predominantly (if not solely) on import to maintain adequate supplies of glutathione, despite encoding *gshAB*, a dual functional enzyme with glutamate-cysteine ligase and glutathione synthetase domains (43). While most streptococci do not possess the capacity to synthesize glutathione and appear to strongly rely on GshT to acquire glutathione, *S. agalactiae* and *S. mutans* are among the few streptococcal species encoding GshAB (43,44). While GshAB is responsible for the bulk of intracellular glutathione in *S. agalactiae* and linked to virulence in a mouse model (44), the *S. mutans* GshAB has been poorly characterized other than one study showing that *gshAB* inactivation impacts the competitive fitness of *S. mutans* against peroxigenic *S. sanguinis* (43), and investigations that support our findings by showing that *S. mutans* cannot synthesize glutathione in a glutathione-free rich media (42,45). Although not a primary objective of this study, our findings underscore the importance of further elucidating the distinct contributions of GshAB and GshT to the maintenance of glutathione homeostasis and their respective roles in the pathophysiology of *S. mutans* and other streptococcal species.

Even though glutathione supplementation did not significantly improve *S. mutans* tolerance to zinc, plate titration assays revealed that the “glutathione-free” Δ*gshT* strain was slightly more sensitive to zinc stress when compared to the parent strain, supporting the idea that glutathione plays a minor role in zinc tolerance. This relatively minor role was within the expected as it has been previously show that the absence of intracellular glutathione had no effect on the zinc tolerance of *E. coli* unless both glutathione import and zinc export genes were simultaneously inactivated (27). Among the other metals tested, cadmium tolerance was also reduced in Δ*gshT* with differences in cobalt sensitivity being marginal but noticeable in the Δ*zccE*Δ*gshT* strain. Cadmium sensitivity was greatly increased in Δ*zccE*Δ*gshT* compared to Δ*zccE*, which is consistent with cadmium’s high affinity for thiol groups (28,46). In eukaryotes, the formation of GSH-metal complex for heavy metal efflux via ABC transporters has been documented, and at least one instance of this mechanism has been reported in bacteria (*Novosphingobium aromaticivorans*) with the formation of a Hg(GS)_2_ complex (41). While it does not appear to be the case in zinc tolerance, it is possible that that the role of glutathione in bacterial metal tolerance may extend beyond intracellular metal sequestration/buffering. Collectively, these results support the role of glutathione as an auxiliary mechanism for bacterial metal tolerance albeit the magnitude and specificity of this protective effect vary among bacteria as well as the affinity and stability of the GSH:metal complex that is formed.

The most unexpected outcome of this investigation was the observation that, contrary to glutathione’s generally beneficial role, upregulation of glutathione uptake genes in the Δ*zccE* strain led to massive accumulation of glutathione that was detrimental to the cell when combined to excess zinc. At this point, we considered two non-mutually exclusive explanations for this phenotype. First, it is conceivable that dysregulated glutathione uptake alone caused the growth inhibition, as supplementation of the medium with high concentrations of glutathione (200 μM or more) led to modest reductions in final growth yields. The second, and most plausible explanation is that rather than conferring protection against zinc intoxication, GshT-mediated uptake of glutathione functions as a “Trojan horse” facilitating zinc entry through the import of GSH:Zn complexes. While the negative effect of the excessive glutathione accumulation on Δ*zccE* growth should not be disregarded, quantification of intracellular glutathione and zinc pools strongly supported the later possibility.

In conclusion, here we provide supporting evidence that glutathione contributes to intracellular zinc homeostasis through formation of GSH:Zn^2+^ coordination complexes that protect cells from zinc-induced mismetallation. Accidentally, we also found that, to manage extremely high intracellular zinc levels caused by the loss of ZccE, *S. mutans* increases glutathione uptake, inadvertently promoting zinc influx through import of GSH:Zn complexes. This later observation also indicates that extracellular GSH:Zn complexes may serve as an alternative mechanism for zinc acquisition in bacteria, independent of the canonical zinc transporter AdcABC.

## ACKNOWLEDGEMENTS

We wish to thank the University of Florida Mass Spectrometry Research and Education Center for their assistance in glutathione quantification via Mass Spectometry, and the University of Florida Center for Environmental and Human Toxicology for ICP-MS analysis.

