## Supplemental Information for "Zinc Tolerance Through Glutathione Import Strikes a Fine Balance Between Protection and Damage in *Streptococcus mutans*"

### Supplemental Material

**TABLE S1.** Primers used in this study

| Primer | Sequence (5' – 3') | Application |
|---|---|---|
| **1942c.del.3arm.rev** | AGGTTCATCAACGCCTTTGGC | *gshT* deletion |
| **1942c.del.3arm.fwd** | CCTTGGAGGATCCAGCTATCTTCCAG | *gshT* deletion |
| **1942c.5arm.rev.del** | GATTTGTTGTGGATCCAACAGTCGCAA | *gshT* deletion |
| **1942c.5arm.fwd.del** | GCGATCATCGTGGTGCTTTAGTAG | *gshT* deletion |
| **1942c.3arm.screen** | GTCATGGTCATCACACCAACAGTTAAC | *gshT* deletion screening |
| **1942c.5arm.screen** | CCCATCCTCTTTCTTTATGACGACAAG | *gshT* deletion screening |
| **zccE.checkF** | GTATTATCAACAACGTATTGCA | *zccE* deletion screening |
| **zccE.checkR** | TTATTGCTTACGAAGATATCG | *zccE* deletion screening |
| **gshTcomp.GA.erm.fwd** | ATATTTTACTGGATGAATTGTTTTAGTAGACTTAACAATG ATATTCGTGATAG | Creation of $gshT^{comp}$ |
| **gshTcomp.GA.erm.rev** | GCCATTTATTATTTCCTTCCTCTTTTACTCAGATTAGACT AGGTGAC | Creation of $gshT^{comp}$ |
| **gshT.qrt.fwd** | GTTCCAACAGTCGCAAGGGT | qRT-PCR of *gshT* |
| **gshT.qrt.rev** | GAGCGGTTGCTTTTGCCTCA | qRT-PCR of *gshT* |
| **tcyA.qrt.fwd** | ATTCAGTTGGCGGGCAGTCT | qRT-PCR of *tcyA* |
| **tcyA.qrt.rev** | CAAGATCGGCGTCCCCCTTA | qRT-PCR of *tcyA* |
| **gyrA.qrt.fwd** | CTCCTGACAAGCCGCATAAA | qRT-PCR of *gyrA* |
| **gyrA.qrt.rev** | GCTCATACGCGCTTCTGTATAA | qRT-PCR of *gyrA* |

**Table S2.** R packages used in data analysis and visualization

| R package | Citation |
|---|---|
| tidyverse | Wickham et al. 2019 (1) |
| ggtext | Wilke & Wiernik, 2022 (2) |
| rstatix | Kassambara, 2023 (3) |
| dbplyr | Wickham et al., 2026 (4) |
| ggpubr | Kassambara, 2023 (5) |
| ggsignif | Ahlmann-Eltze & Patil, 2021 (6) |
| multcomp | Hothorn et al., 2008 (7) |
| magrittr | Bache & Wickham, 2022 (8) |
| emmeans | Lenth et al., 2025 (9) |
| lme4 | Bates et al., 2015 (10) |

**Fig S1**

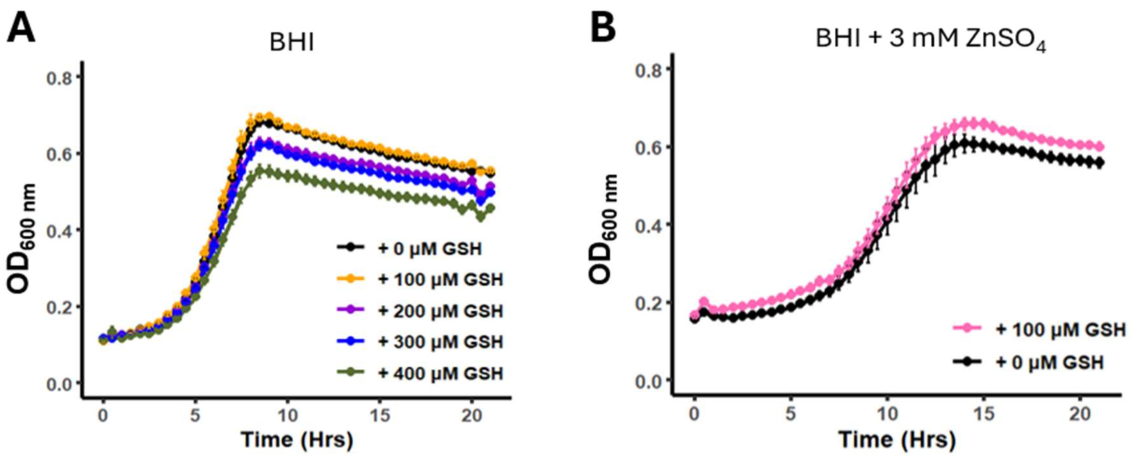


**Fig. S1.** Glutathione supplementation minimally improves *S. mutans* UA159 growth on a sub-inhibitory concentration of zinc. (A) Growth of UA159 in BHI supplemented with increasing concentrations of glutathione (GSH). (B) Growth of UA159 in BHI + 3 mM $ZnSO_4$ with or without 100 μM glutathione (GSH) supplementation. Error bars are represented by SEM.

### Fig S2

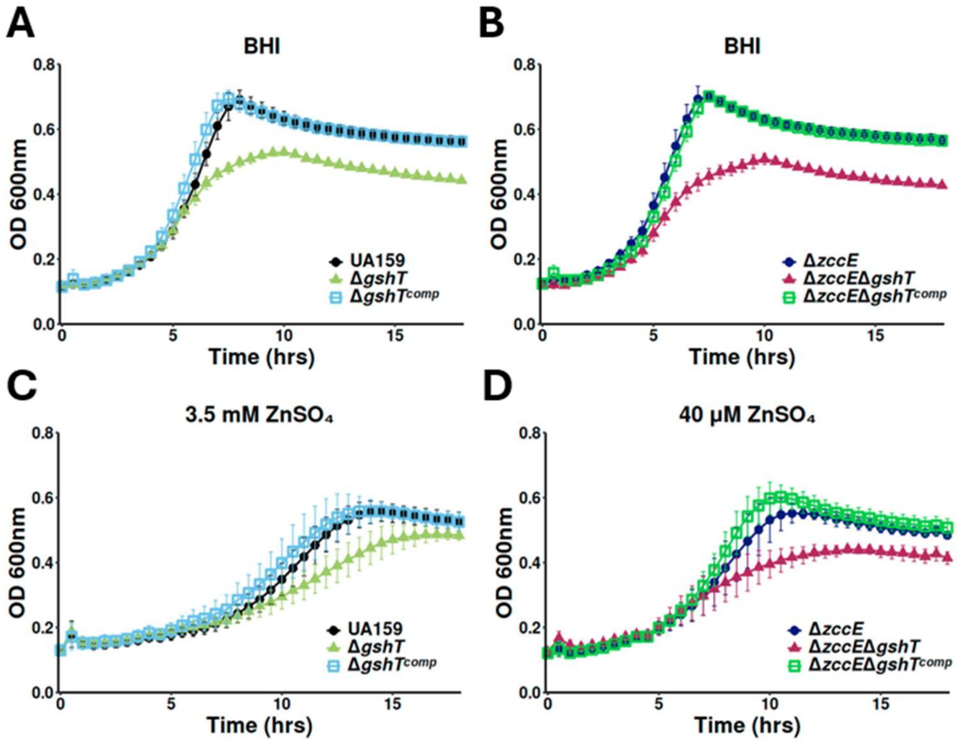


**Fig. S2.** Growth of *S. mutans* UA159, Δ*gshT*, Δ*gshT*$^{comp}$, Δ*zccE*, Δ*zccE*Δ*gshT* and Δ*zccE*Δ*gshT*$^{comp}$ in plain BHI (A-B) or BHI + 3.5 mM $ZnSO_4$ (C) or BHI + 40 μM $ZnSO_4$ (D).
